# Ensemble–function relationships to evaluate catalysis in the ketosteroid isomerase oxyanion hole

**DOI:** 10.1101/2021.09.29.461692

**Authors:** Filip Yabukarski, Tzanko Doukov, Margaux Pinney, Justin Biel, James Fraser, Daniel Herschlag

## Abstract

Following decades of insights from structure–function studies, there is now a need to progress from a static to dynamic view of enzymes. Comparison of prior cryo X-ray structures suggested that deleterious effects from ketosteroid isomerase (KSI) mutants arise from misalignment of the oxyanion hole catalytic residue, Y16. However, multi-conformer models from room temperature X-ray diffraction revealed an ensemble of Y16 conformers indistinguishable from WT for Y32F/Y57F KSI and a distinct, non-native ensemble for Y16 in Y57F KSI. Functional analyses suggested rate effects arise from weakened hydrogen bonding, due to disruption of the Y16/Y57/Y32 hydrogen bond network, and repositioning of the general base. In general, catalytic changes can be deconvoluted into effects on the probability of occupying a state (*P*-effects) and the reactivity of each state (*k*-effects). Our results underscore the need for ensemble–function analysis to decipher enzyme function and ultimately manipulate their extraordinary capabilities.

## MAIN TEXT

Structure–function investigations have been central to our understanding of macromolecular function and enzyme catalysis (e.g., (*1, 2*)). For enzymes, tens of thousands of crystal structures from cryogenic (cryo, ∼100 K) X-ray diffraction data have shown that catalytic and reactant groups are positioned in enzyme active sites to interact with substrates and transition state analogs and facilitate reactions. Supporting the importance of positioning, mutations that misposition catalytic groups are often associated with reduced catalysis (e.g., (*3*–*6*)). Nevertheless, atoms in proteins are constantly in motion, so their positions vary. Thus, proteins do not consist of a single conformational state but of an ensemble of conformational states defined by an energy landscape (*7*–*12*); cryo X-ray structures are snapshots of this larger ensemble (*13*–*16*). Accordingly, an observed reaction rate is the aggregation of the reaction from each conformer on the landscape, with different conformers having different contributions dependent on their abundance and reactivity. Thus, ensemble information is required to relate structure to conformational landscapes and to relate structural states to functional properties.

Recognizing these limitations, we previously obtained information about the conformational ensemble of the model enzyme ketosteroid isomerase (KSI) in its apo state and with bound reactants and transition state analogs, allowing general and specific catalytic proposals to be tested (*17*). Here we probe the conformational ensembles of two KSI mutants and apply “ensemble–function” analysis to decipher how mutations affect function. We first demonstrate the limitations of traditional structure–function approaches and how expanding from static structure into the realm of conformational ensembles can overcome these limitations. We then describe KSI ensemble–function experiments and analyses that allow us to test models that were previously indistinguishable and provide evidence for previously unrecognized catalytic behaviors.

### KSI structure–function and the need for ensemble–function

Ketosteroid isomerase (KSI) binds its steroid substrates in a hydrophobic pocket and, for catalysis, uses an oxyanion hole, consisting of hydrogen bond donors Y16 and D103, and a general base, D40, that shuffles protons in steroid substrates (**Figure 1A, B**; (*18, 19*)). Site-directed mutagenesis studies reveal large deleterious rate effects from removal of these side chains as well as effects from mutations of groups interacting with them (*20*–*23*).

**Figure 1.**
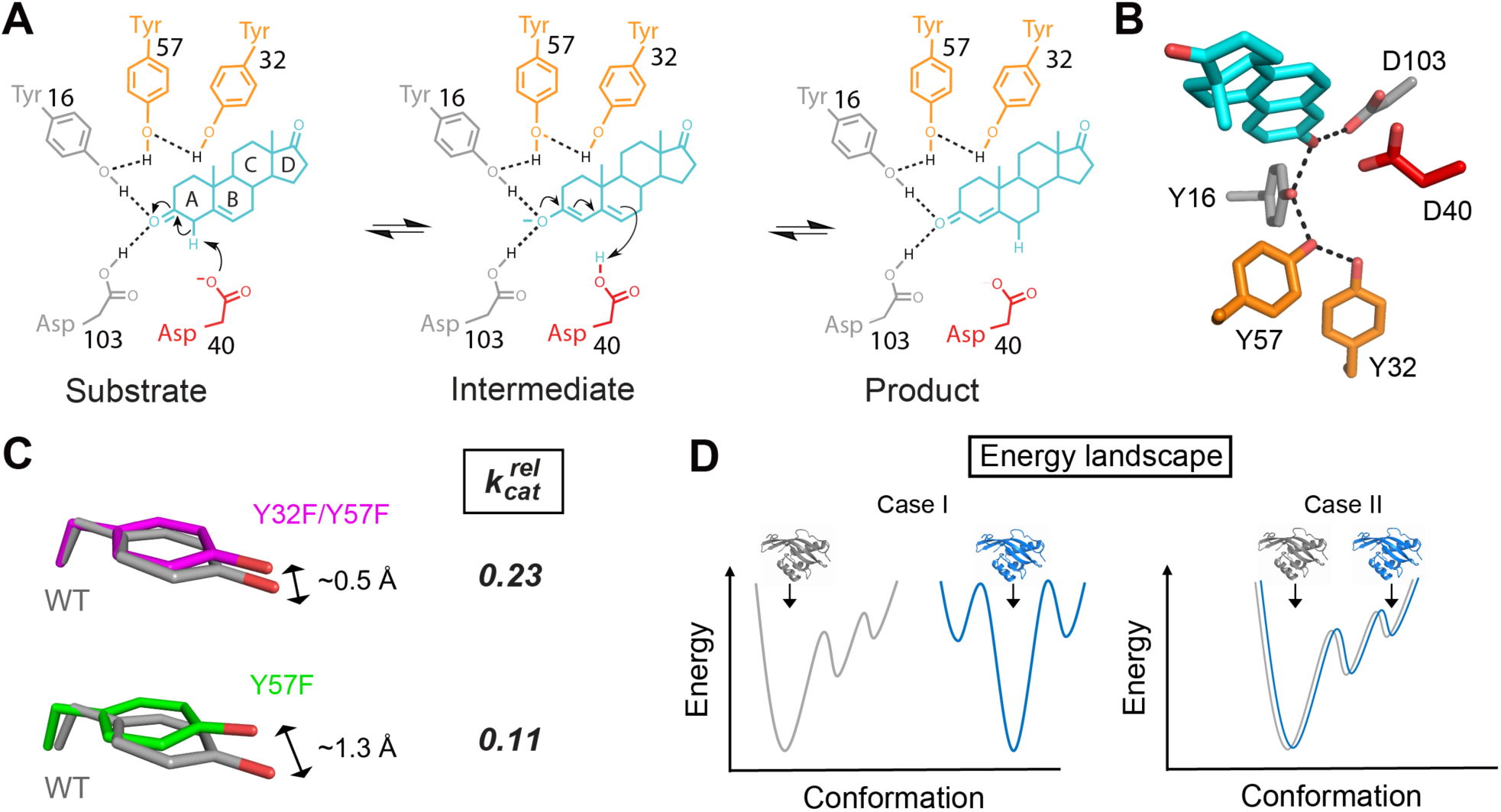
KSI reaction mechanism, active site, and structure-function results. (**A**) KSI catalyzes double bond isomerization of steroid substrates (shown for the substrate 5-androstene-3,17-dione) utilizing a general acid/base D40 (which we refer to herein as a general base), and an oxyanion hole composed of the side chains of Y16 and D103 (protonated). (**A, B**) Y16 is embedded within a hydrogen bond network with two other tyrosine residues, Y57 and Y32. The general base, oxyanion hole, and hydrogen bond network residues are colored in red, grey, and orange, respectively. Structural model 1OH0 from the PDB (*18*). (**C**) In Y32F/Y57F KSI (PDB 1DMN (*3*)) and Y57F KSI (PDB 1DMN (*3*)), Y16 (magenta and green, respectively) is misaligned with respect to its position in WT KSI (grey, PDB 3VSY (*24*)) (see **Table S1** and Materials and Methods for alignment RMSDs and procedures, respectively; also see **Figure S1**). The mutant *k*_cat_ values are shown relative to WT ((*3*), also see **Table S2**). (**D**) An observed structural difference between WT and a mutant can arise either because the underlying conformational ensembles of the molecules are different (Case I) or because conditions trapped different states in the cryo-cooled structures from a common ensemble (Case II). In Case I the grey (left) and blue (right) conformational landscapes are distinct and the crystal structures have captured distinct states from each ensemble (indicated by arrows), whereas in Case II the grey and blue conformational landscapes are identical, but the crystal structures have captured distinct states (indicated by arrows).

Prior structure–function studies revealed mutations in the KSI Y16 hydrogen bond network result in ∼0.5 Å (Y57F/Y32F KSI) and ∼1.3 Å (Y57F KSI) displacements of Y16 from its WT position (**Figure 1C** and **Figure S1**) (*3*). As Y16 donates a hydrogen bond that stabilizes the anionic transition state (**Figure 1A**), the simplest interpretation of these results is that these displacements are responsible for the observed 4- and 9-fold decreases in catalysis, with the larger displacement resulting in the larger rate decrease (**Figure 1C** and **Table S2**). However, as elaborated and demonstrated below, ensemble information and deeper functional analyses are needed to evaluate these effects and uncover their physical origins.

There are fundamental limitations with the traditional structure-function approach taken above. First, individual X-ray structures do not provide conformational ensembles (*9, 10, 25*). Instead, individual X-ray structures obtained at cryo temperatures can capture different states within an ensemble (*13*–*16*). In addition, cryo temperatures can alter the ensembles of states and thus not accurately capture the states occupied or predominant in the ensemble present at physiological temperature (*26*–*29*). While crystallographic B-factors provide some information that could be related to ensemble properties, B-factors include model uncertainties and errors, are incomplete models for heterogeneity, and cannot be used to determine the ensemble of conformational states present from a single cryo X-ray structure (*30, 31*). Further, as different conformational states can be observed in individual cryo-cooled crystals of the same protein (*14*–*16*), differences between cryo X-ray structures of WT and mutant variants can arise either because the underlying conformational ensembles of the molecules are different or because different conditions trapped different states in the cryo-cooled structures from a common ensemble (**Figure 1D**). Ensemble information is required to distinguish between these cases and to draw conclusions about the presence and extent of conformational changes.

The second important limitation of traditional structure–function correlations is that even if an observed conformational difference truly reflects different ensembles for the enzymes under comparison, this conformational difference may or may not be responsible for the observed rate effect. Additional functional studies are required to test whether the change is causative or simply correlative and to determine the origin of the observed rate effects.

### Structure–function to ensemble–function analysis of Y32F/Y57F KSI

We first consider Y32F/Y57F KSI, which exhibits a 4-fold rate decrease relative to WT KSI (**Figure 1C** and **Table S2**). In the simplest scenario, 1/4^th^ of the Y16 mutant ensemble samples reactive WT conformations and 3/4^th^ samples alternative, non-reactive conformations. A second possibility is that the WT configuration is even less populated, and the reaction is 4-fold slower in the alternative conformational state (**Figure 2A**). Both models (and models between these extremes) presuppose a difference in the WT and mutant conformational ensembles.

**Figure 2.**
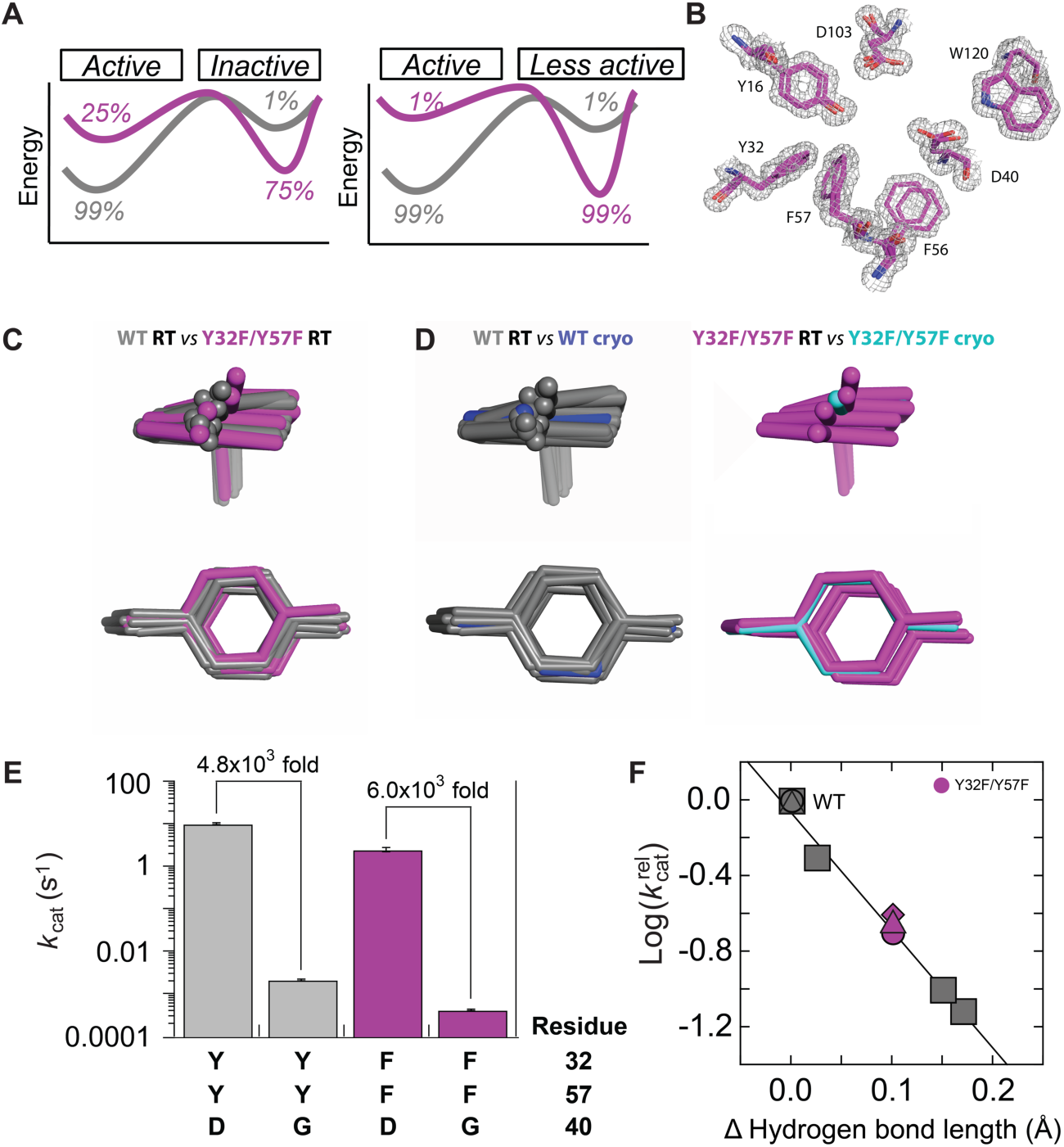
Ensemble and functional data for Y32F/Y57F and WT KSI. **A**. Ensemble models for the 4-fold effect in Y32F/Y57F mutant. Left: in Y32F/Y57F KSI, Y16 samples reactive WT conformations 1/4^th^ of the time (25%) and alternative non-reactive conformations 3/4^th^ of the time (75%), whereas WT KSI is predominantly in reactive conformations (99%). Right: in Y32F/Y57F KSI, reactive WT conformations are populated by Y16 less than one-fourth of the time (1% in this schematic), and Y16 predominantly populates less reactive alternative conformations that are responsible for the observed reaction. **B**. Representative electron density (grey mesh) and multi-conformer modeling (magenta sticks) for the Y32F/Y57F KSI active site. Electron density is contoured at 1 σ. **C**. Overlay of the WT (grey sticks; (*17*)) and the Y32F/Y57F ensembles (see Materials and Methods and **Table S4** for alignment RMSDs). **D**. Superposition of the cryo crystal structures with the RT ensemble models suggests that the cryo structures have captured Y16 conformations from a common ensemble of states. Left: WT RT ensemble (grey, (*17*)) and the WT cryo structure (PDB 3VSY (*24*)). Right: Y32F/Y57F ensemble (magenta) and the Y32F/Y57F cryo structure (PDB 1DMN, (*3*)). **E**. The same rate effect is observed from ablating the general base D40 in WT (grey) and in Y32F/Y57F (magenta) for reaction of 5(10)-estrene-3,17-dione (see **Figure S4** and **Table S2**). **F**. Catalytic effects in KSI variants *versus* changes in the hydrogen bond length with a bound TSA for KSI variants. Grey squares reproduce data from Pinney et al. (*37*) and give a correlation with R^2^ = 0.99. Y32F/Y57F kinetics relative to WT (magenta triangle) with the substrate 5(10)-estrene-3,17-dione and Y32F/Y57F kinetics relative to WT (magenta diamond) with the substrate 5-androstene-3,17-dione, and Y32F/Y57F/D40G relative to D40G (magenta circle) with the substrate 5(10)-estrene-3,17-dione (see **Table S2** for kinetic constants and **Table S5** for hydrogen bond lengths).

To test whether there is indeed an altered conformational ensemble for the Y32F/Y57F KSI mutant, we collected room temperature X-ray diffraction data (RT X-ray). X-ray data for crystals obtained at temperatures above the glass transition (∼180–220 K) provide information about conformational heterogeneity, which is the experimental manifestation of conformational ensembles (*32*–*36*). We used the 1.10 Å RT X-ray data to obtain Y32F/Y57F KSI multi-conformer models that capture the conformational heterogeneity in the crystal (**Table S3**; **Figure 2B**, see **Materials and Methods**). We then compared the Y32F/Y57F with the WT ensemble obtained previously (*17*) and observed that the Y16 states extend in the same directions and span the same range (**Figure 2C**). Comparison to the cryo X-ray models suggests that the different cryo structures for WT and Y32F/Y57F KSI represent individual states within a common ensemble (**Figure 2D**).

Given the highly similar ensemble of Y16 states for WT and Y32F/Y57F KSI, we needed to consider alternative models to account for the rate difference between these variants. These models fall into two classes: a direct effect on the Y16 hydrogen bond or indirect effects on other catalytic elements. There was no significant change in the ensemble of the other oxyanion hole hydrogen bond donor, D103 (**Figure S2**), in substrate binding (**Table S2**), and mutational ablation of the general base (D40G) gave the same rate reduction for WT and Y32F/Y57F KSI (**Figure 2E** and **Table S2**), providing no indication of effects on other catalytic features. We therefore turned to consideration of energetic effects within the oxyanion hole.

Prior results established a linear free energy relationship (LFER) between the length of KSI oxyanion hydrogen bond donors and the amount of catalysis (**Figure 2F**, grey points (*37*)). Specifically, mutations that alter the partial positive charge on the oxyanion hole hydrogen bond donors, such as D103N, give a systematic and linear relationship between how much the hydrogen bond is lengthened and how much catalysis reduced; a longer hydrogen bond is weaker and provides less stabilization for the oxyanionic transition state, relative to the carbonyl ground state (or water-occupied site in the apo enzyme) (*38*). This relationship is distinct from proposals of special energetic contributions from short or symmetrical hydrogen bonds (see **supplementary text 1**).

^1^H NMR chemical shifts have been shown to report on changes in hydrogen bond length (*38*–*41*). In Y32F/Y57F KSI, we observed that the ^1^H NMR chemical shift of the Y16-oxyanion hydrogen bond proton shifted upfield, indicating a lengthening of the Y16 hydrogen bond by 0.1 Å, relative to WT KSI ((*37*), **Table S5**). Applying this lengthening to the above-noted LFER predicts the observed 4-fold rate effect (**Figure 2F**, magenta points). This lengthening is expected on chemical grounds for the Y32F/Y57F mutation, as removal of the neighboring hydrogen bond donor will lessen polarization of the Y16 hydroxyl group and thereby yield a longer and weaker hydrogen bond (**Figure 4A, B**) ((*42, 43*); also see **supplementary text 2**)

In summary, the initial comparison of individual cryo X-ray structures of WT and Y32F/Y57F KSI was misleading, as it reflected single structures randomly selected from each ensemble. Multi-conformer models from RT X-ray data revealed a highly similar ensemble of conformational states, and further analysis of the Y16 oxyanion hole hydrogen bond revealed a lengthening and weakening, presumably due to loss of the Y57/Y32 hydrogen bond network that polarizes Y16 (**Figure 4A, B**). The weakened hydrogen bond is predicted to decrease catalysis by 4-fold, as is observed. A combination of ensemble and functional data (ensemble–function analysis) allowed us to evaluate and distinguish catalytic models.

### Ensemble–function analysis of Y57F KSI

We now turn to the Y57F mutation in KSI, where analogous ensemble and functional experiments revealed a more complex scenario than that for Y32F/Y57F KSI, but one that could nevertheless be disentangled through ensemble–function studies. Given the 9-fold rate decrease relative to WT KSI, in the simplest scenario, Y16 in this mutant would sample reactive WT conformations about 10% of the time and would spend the remaining 90% of the time in alternative, non-reactive conformations (**Figure 3A**, left). The other ensemble model, as above, is that Y16 in the Y57F mutant samples alternative, less reactive conformations and reacts from those while sampling WT conformations less than 10% of the time (**Figure 3A**, right).

**Figure 3.**
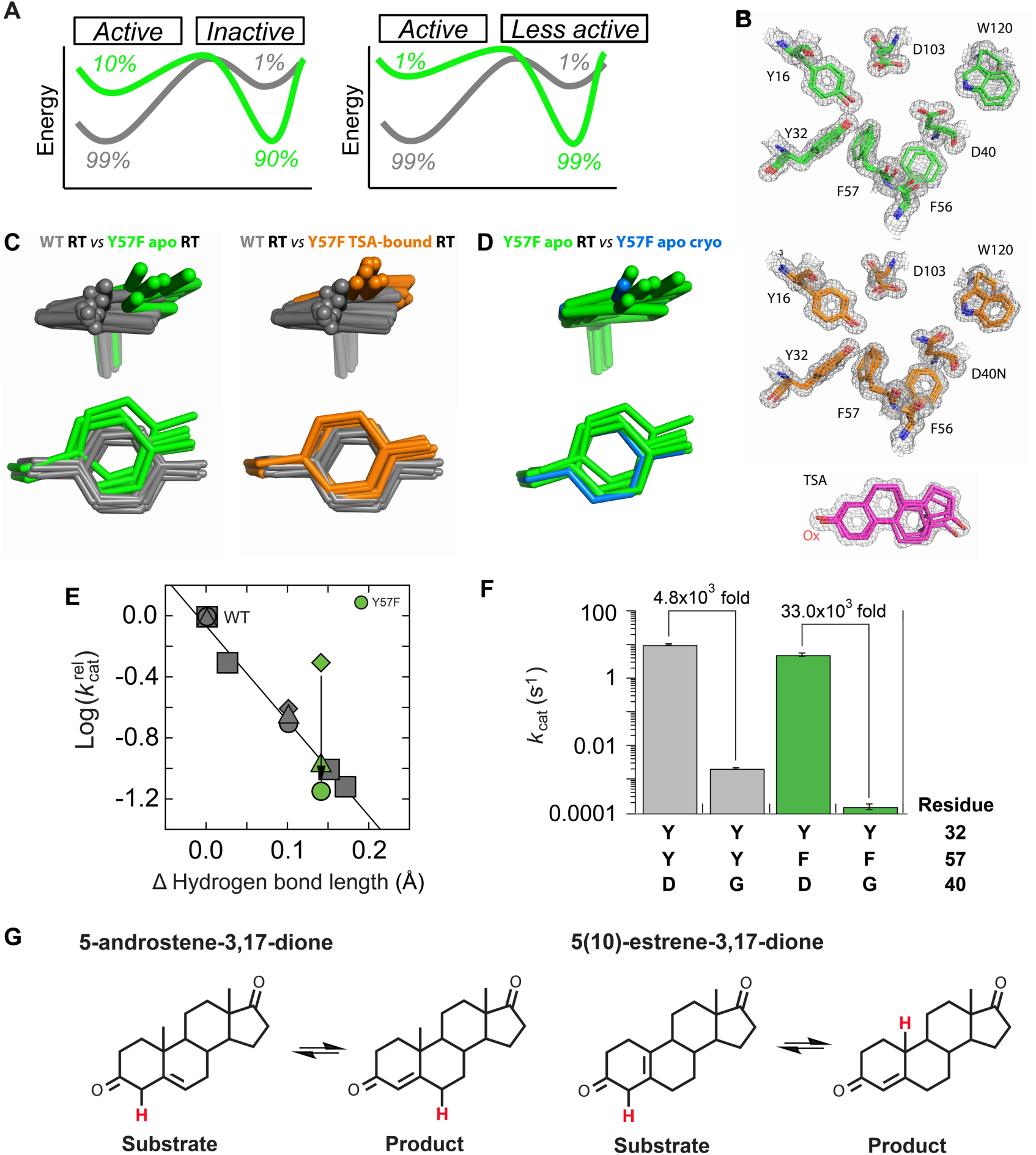
Ensemble and functional data for Y57F and WT KSI. **A**. Ensemble models for the 9-fold effect in Y57F mutant. Left: in Y57F Y16 samples reactive WT conformations ∼1/10^th^ of the time (∼10%), while spending ∼9/10^th^ of the time (∼90%) in alternative, non-reactive conformations. Right: in Y57F KSI, reactive WT conformations are not significantly populated by Y16 which instead reacts (less efficiently) from alternative conformations that it predominantly populates. **B**. Representative electron density (grey mesh) and multi-conformer modeling for the KSI Y57F apo (top, green sticks) and Y57F (D40N) TSA-bound (bottom, orange sticks) active site. Also shown is the stick model (pink) and associated electron density (grey mesh) of the bound TSA. D40N mutation was introduced to mimic the protonated general base and increase TSA affinity (*44, 45*). Electron density is contoured at 1 σ. **C**. Overlay of the WT (grey sticks, (*17*)) and the Y57F apo (left, green sticks) and Y57F(D40N) TSA-bound (right, orange sticks) ensembles (see Materials and Methods and **Table S4** for alignments). **D** Superposition of the cryo crystal structures with the RT ensemble models suggests that the cryo structure captures a Y16 conformation from a common ensemble of states. Y57F RT ensemble model (green) and the Y57F cryo structure (PDB 1DMM (*3*)). **E**. Plot of catalytic effects in KSI variants *versus* changes in the hydrogen bond length from Figure 2A but now also including Y32F/Y57F data points (grey symbols). Y57F kinetics with the substrate 5-androstene-3,17-dione relative to WT (green triangle), Y57F kinetics relative to WT (green diamond) and Y57F/D40G relative to D40G (green circle) with the substrate 5(10)-estrene-3,17-dione (see **Table S2** for kinetic constants and **Table S5** for hydrogen bond lengths). **F**. Different rate effects from ablating the general base D40 in WT (grey) and in Y57F (green) for reaction of the substrate 5(10)-estrene-3,17-dione (**Table S2**). **G**. KSI reaction with the steroid substrates 5-androstene-3,17-dione (left) and 5(10)-estrene-3,17-dione (right). The shuffled proton is colored in red.

**Figure 4.**
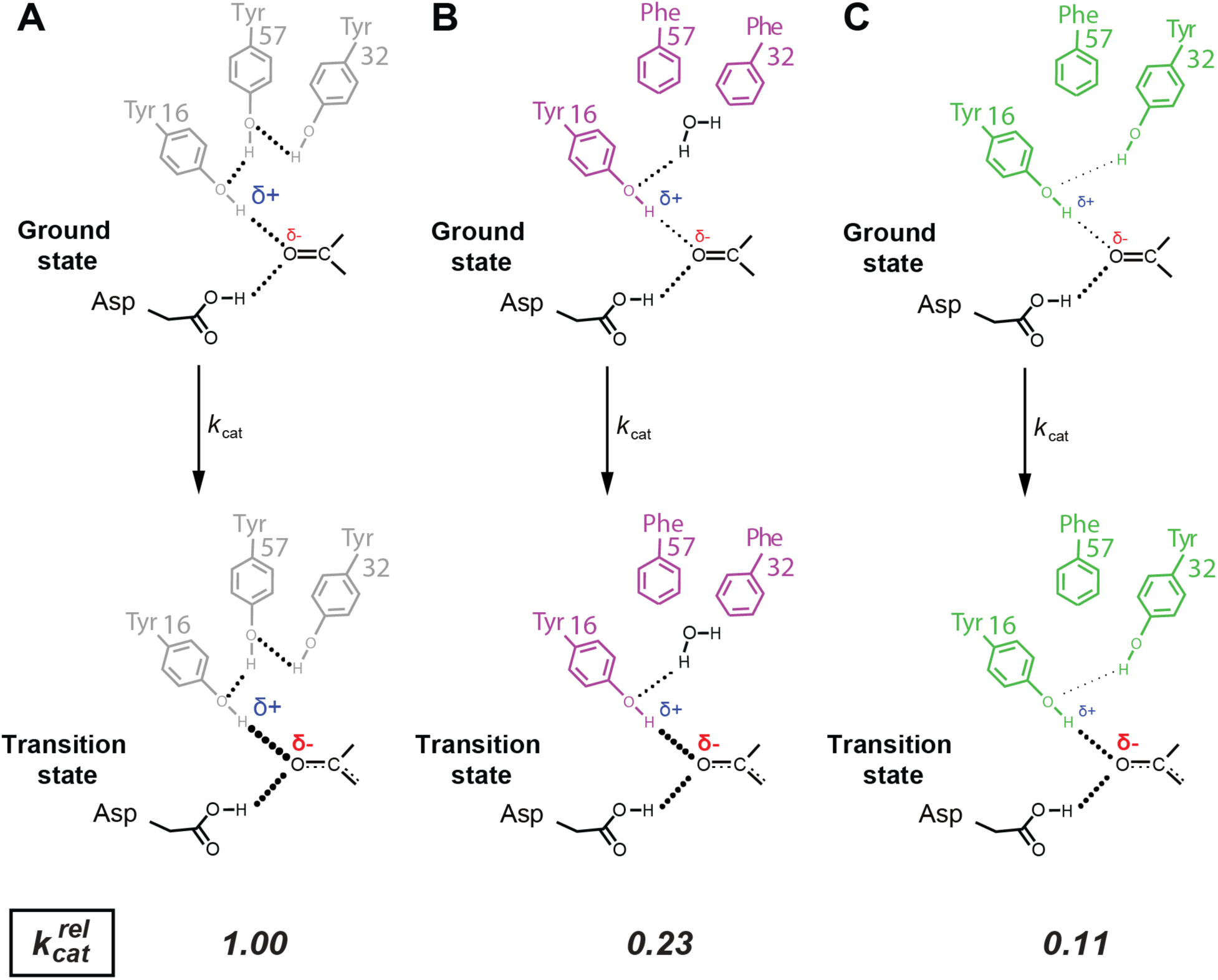
KSI oxyanion hole catalytic model. During the KSI reaction, the amount of negative charge on the substrate carbonyl increases and this negative charge accumulation is stabilized by hydrogen bonds. Analogously, hydrogen bonds become stronger as the charge density on the hydrogen bond donating hydrogen increases (*38, 47*– *49*). Thus, WT (grey), Y32F/Y57F (magenta), and Y57F (green) have decreasing hydrogen charge densities, respectively, and provide lesser extent of transition state stabilization (hydrogen bond strength is depicted by the size of the dots representing the hydrogen bonds). In all cases, hydrogen bonds shorten and strengthen in the transition state (indicated with thicker doted lines in the TS compared to GS), but the shortening and strengthening in the TS decreases in the following order WT, Y32F/Y57F, Y57F.

We examined the conformational ensemble for Y57F KSI with 1.16 Å RT X-ray data and multi-conformer modeling (**Table S3, Figure 3B**). The Y16 ensemble for this mutant is largely distinct from WT KSI with no observed overlap (**Figure 3C**, left, **Figure S2**). This observation is not consistent with the simplest model of fractional occupancy of the WT state (**Figure 3A**, left), and additional evidence counter to this model comes from the set of conformations for Y57F KSI with a bound oxyanion transition state analog, obtained from RT X-ray data at 1.11 Å resolution (**Figure 3B**, bottom, **Table S3**). With bound transition state analog, the positions of Y16 overlap with those for apo Y57F KSI conformational states and remain distinct from those for WT KSI (**Figure 3C**, right, also see **Figure S2**). These data support a model in which the Y57 KSI reaction occurs predominantly via a distinct set of oxyanion Y16 hole conformational states.

Without additional functional data, we might draw the conclusion that misalignment of the Y16 ensemble *causes* the 9-fold catalytic effect from Y57F mutation (**Figure 3A**, right). But we would not know the mechanistic origin of the effect—why the alternatively aligned states are less reactive—or even if the misalignment is correlated or causative. Possible models involve altered substrate binding, weakened oxyanion hole hydrogen bonding, and altered general base catalysis. Recognizing these possibilities, we tested each model.

Remarkably, the 9-fold rate effect observed for reaction of Y57F KSI with the substrate 5-androstene-3,17-dione is predicted from the increased length of the Y16 oxyanion hole hydrogen bond (**Figure 3E**, green triangle), just like the 4-fold effect for Y32F/Y57F KSI (**Figure 2F**, magenta triangle). The loss of the neighboring Y57 hydrogen bond would be expected to lessen polarization of the Y16 hydroxyl group and thereby weaken its oxyanion hole hydrogen bond, as described above (**Figure 4A, C**, also see **supplementary text 1, 2**). We speculate that the longer hydrogen bond and larger effect for the Y57F mutant than for the Y32F/Y57F double mutant arises because solvent access is greater for the double mutant and interactions with solvent water molecules are more effective at polarizing Y16 than the distorted Y16/Y32 hydrogen bond in Y57F KSI (**Table S2**).

These results suggest that the reaction is slower because the Y16 hydrogen bond is weaker, and not because the altered Y16 conformational states in the Y57F mutant give slower reactions (**Figure 4**). A range of conformational states are present within WT KSI’s active site that appear to allow proton transfer at sites that are several Ångstrøms apart, as needed in the KSI reaction (**Figure 1A, Figure 3G**, and **Figure S4**; (*17*)). This range of conformations also appear to allow reaction from the altered conformational poses present in Y57F KSI without further sacrificing catalysis.

In contrast to the observation of the predicted 9-fold effect for Y57F KSI, as described above, a rate effect of only 2-fold was observed for this mutant when reacting with the substrate 5(10)-estrene-3,17-dione (**Figure 3E**, green diamond, **Figure 3G**). As the catalytic effect was less than predicted, we reasoned that there might be a second, compensating effect, such as an increased catalytic contribution from the general base in the Y57F mutant. This model predicts that the Y57F mutation will have a larger catalytic effect with the general base ablated—i.e., in a D40G background—and that that effect would match the 9-fold effect predicted from the LFER in **Figure 3E**. Both predictions are met. The D40G mutation gives a 33×10^3^-fold effect in the Y57F background but only a 5×10^3^-fold effect in WT KSI (**Figure 3F**), and in the D40G background, the Y57F mutation causes a 14-fold rate decrease, within 2-fold of the 9-fold effect predicted by the LFER (**Figure 3E, Table S2**). The altered Y16 ensemble in the Y57F mutant presumably differentially alters the reactivity of the two substrates, which have different geometries and protons that are shuffled between different positions (**Figure 3G** and **Figure S4**). The ensemble of states for the general base (D40) shows no apparent change, so that the reactivity difference may arise from differential placement of the substrate (**Figure S5**).

In contrast, the similar substrate affinities for WT and Y57F KSI (**Table S2**) and prior observation that the B, C and D rings of steroid substrates contribute solely to binding, but not catalysis, once bound provide no indication of effects coupled to the substrate binding site (*46*).

In summary, the Y57F KSI mutation results in a change in the bound and likely in the reactive ensemble. Nevertheless, our observations suggest a model in which the observed 9-fold rate decrease does not arise from these altered states but rather arise from a weakened hydrogen bond. With a different substrate, the reactivity is higher than predicted based on the active site hydrogen bond, and double mutant cycle analyses trace this effect to a fortuitous increase in reactive alignment with general base for this substrate in Y57F KSI. Again, ensemble–function analysis was needed to evaluate and distinguish between catalytic models.

### Implications

The ensemble–function analyses carried out in this study revealed a very different, and far richer, mechanistic landscape than accessible via traditional structure–function analyses. Consider the hypothetical observation of a mutation that gives a deleterious rate effect and structural change. The simplest and most common interpretation would be that the observed change in structure *causes* the change in catalysis. However, multiple models are possible, as we describe for two KSI mutants, and we distinguished these models by integrating structural data that report on conformational ensembles, rather than static structures, with synergistic functional studies.

In general, the observation of an altered conformation from comparison of cryogenic X-ray structures for a WT and mutant proteins does not indicate that the ensemble has in fact changed; cryo-freezing of crystals traps a protein in different conformational poses that correspond to states on a conformational landscape, so common or different structures can arise when WT and mutant landscapes are identical or overlapping ((*14*–*16*), **Figure 1E**). Indeed, we observed an apparent change for Y32F/Y57F KSI based on individual cryo X-ray structures, but room temperature X-ray data revealed similar, overlapping conformational ensembles for this mutant and WT KSI (**Figure 2**).

Functional effects can arise from alterations in the occupancy of states that differ in reactivity or from changes in the reactivity of the individual states, as was the case herein from a weakened oxyanion hole hydrogen bond (**Figure 2F, 3E**). Additional complexities arise as these effects can differ for different substrates. These differences underscore how conformational landscapes can be exploited to evolve enzymes that use new substrates and catalyze new reactions (*50*–*52, 52, 53*) and, conversely, underscore the need to probe function from an ensemble perspective to understand how enzymes have evolved.

Most generally, the rate of an enzyme-catalyzed reaction is a function of the occupancy of each state on a multi-dimensional energy landscape and the probability of reacting from that state (**Figure 5**). Thus, observed functional changes can arise through “*k*-effects” on the binding, rate or other functional properties of each state (**Figure 6A, B**) or through “*P*-effects” on the probability distribution of states (**Figure 6C, D**). *P*-effects can occur in two subtypes, lessening the amount of the most active state but still reacting predominantly through that state (**Figure 6C**) or reacting predominantly through an alternative state that becomes more prevalent (**Figure 6D**).

**Figure 5.**
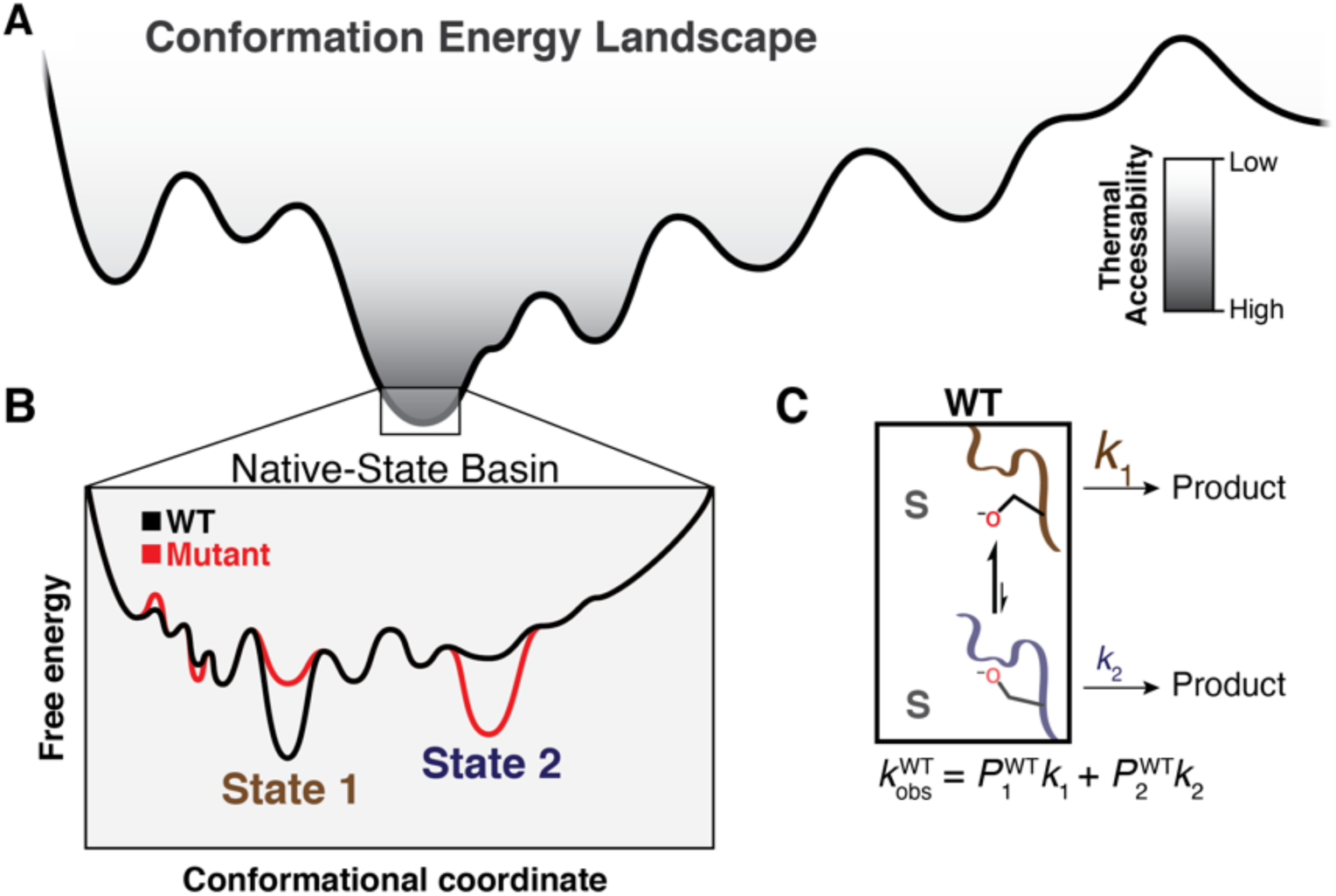
Catalysis from an ensemble perspective. **A**. Enzymes form a set of states specified by energy wells on a free energy landscape, with dimensionality defined by the thousands of degrees of freedom from each rotatable bond of each residue, depicted here schematically in a single dimension. **B**. An example ensemble of near-energy substates in which State 1 (brown) and State 2 (blue) lie within the lowest-energy basin (“Native-State Basin”). These substates have different intrinsic reactivities, reflecting different barrier heights along their individual reaction coordinates (see **Figure 6**). **C**. Mathematically, the observed rate constant of the WT (*k*_obs_^WT^) with substrate S is the probability-weighted (occupancy-weighted) sum of the intrinsic rate constants of each microscopic substate. Here we show a simplified example with two states; this example can be generalized across all states with sufficient occupancy and reactivity to contribute significantly to the observed reaction rate: *k*_*obs*_ = ∑_*i*_ *P*_*i*_ + *k*_*i*_ where *P* is the probability of occupying state i and *k* is the rate of reacting from that state. Modified from (*54*).

**Figure 6.**
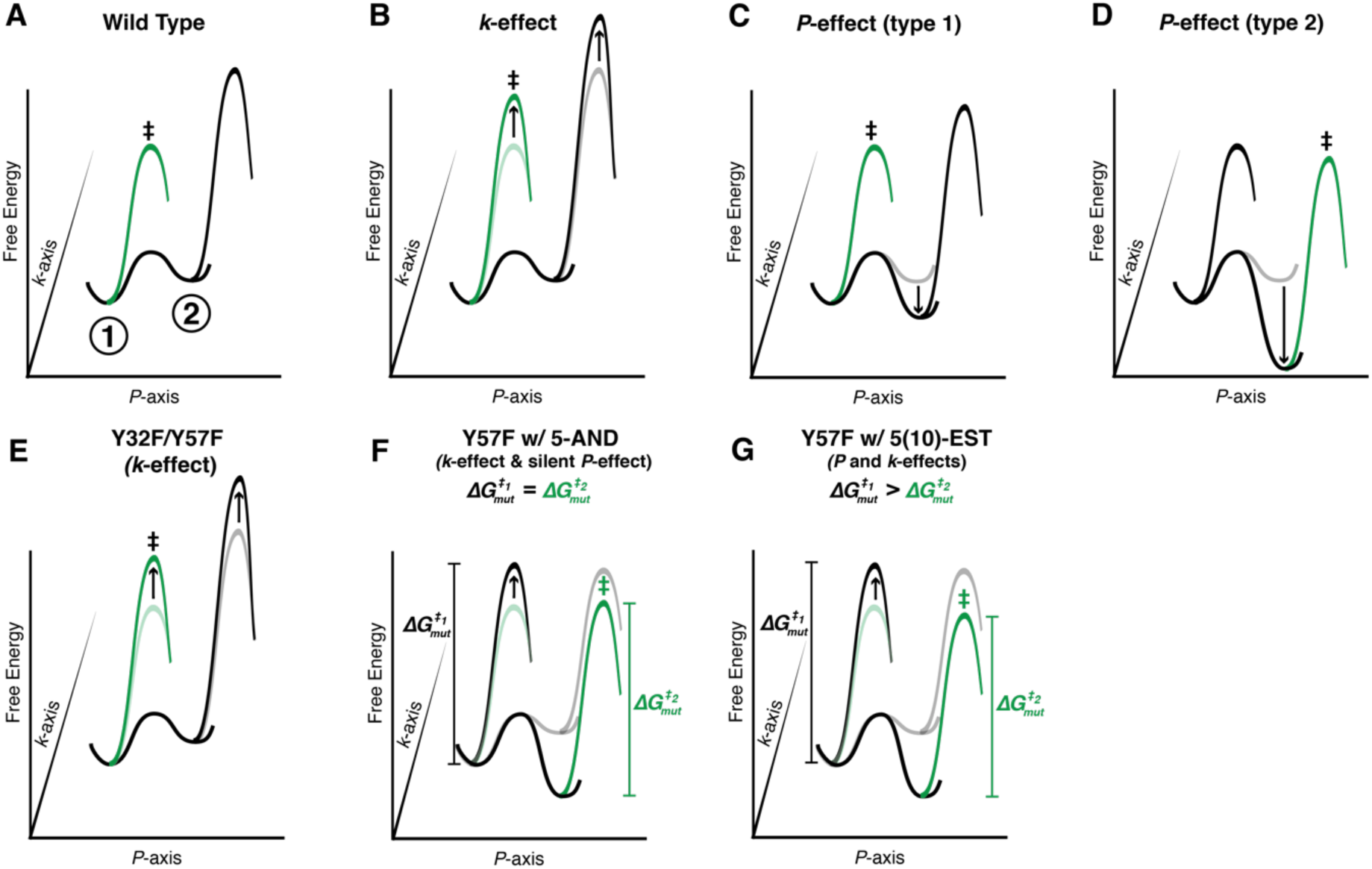
The effects of functional (*k*-effects) and occupancy changes (*P*-effects) to reactivity from an enzyme ensemble. In all panels, the *k*-axis is the reaction coordinate, the *P*-axis is the conformational coordinate, here simplified to two conformational states, and the Z-axis is free energy. Mutant profiles are in grey and light green, and the green profiles (with ≠) represent the preferred reaction path. **A**. A simplified ensemble reaction coordinate for a WT enzyme that reacts preferentially from the most active and most probable state (green). A less reactive and less probable state is also depicted (black). **B**. Depiction of a *k*-effect, which increases the barrier to reaction uniformly in both states and reactions occur via the most populated state (≠, green). **C**. Depiction of a *P*-effect that changes the occupancy of states, but not the most reactive conformation. Reduced reactivity results from decreased occupancy of the more-reactive state. **D**. Depiction of a *P*-effect that results in the enzyme reacting from a more probable, but less reactive conformation (≠, green). **E**. The Y32F/Y57F KSI mutant displays a “*k*-effect” that decreases reaction rate by 4-fold in all conformers. **F**. The Y57F KSI mutant with the substrate 5-androstene-3,17-dione (5AND) displays a *k*-effect that decreases reaction rate by 9-fold in all conformers and a silent *P*-effect where the state probabilities change but this change is not responsible for the observed rate effect. **G**. The rate effect observed for the Y57F KSI mutant with the substrate 5(10)-estrene-3,17-dione (5(10)-EST) displays a combined *P*- and *k*-effect, where the increase in rate is mitigated by an *increase* in occupancy of a state that is better aligned for reaction with this substrate.

**Figure 6E–G** depict the *k*- and *P*-effects for our KSI mutants. The comparison in **Figure 6E** represents the case observed for Y32F/Y57F *vs*. WT KSI, where there is no change in the occupancy of states—no *P*-effect—but there is a weakening of the Y16 oxyanion hole hydrogen bond that causes a uniform increase in the reaction barrier (“*k*-effect”; see also **Figure 6B**). The reaction of Y57F KSI with the substrate 5-androstene-3,17-dione is a special case of the “*P*-effect”, where the state probabilities change but this change is not responsible for the observed rate effect; instead, the observed change in rate is from a *k*-effect (due to the weakened Y16 oxyanion hole hydrogen bond) that is common to all states (**Figure 6F**). Thus, the rate effect simply reflects the extent of hydrogen bond weakening. A more complex scenario arises for reaction of Y57F KSI with the second substrate 5(10)-estrene-3,17-dione, where the observed rate effect arises from a combination of weakening the Y16 hydrogen bond (as above) and mitigation of this effect from *increased* occupancy of states that are better aligned for the different proton transfer needed by this substrate (**Figure 6G**). According to this model, Y57F KSI reactivity is down ∼9-fold due to a weaker hydrogen bond, but increased by 5-fold due to more favorable alignment of the general base, together resulting in the observed overall effect of 2-fold.

These results are consistent with prior ensemble data suggesting that equivalent KSI oxyanion hole hydrogen bonds can be formed across a range of conformational states, and the results herein support the model of KSI oxyanion hole catalysis, relative to reaction in aqueous solution, arises because the oxyanion hydrogen bonds are stronger than those to water, due to the presence of intrinsically stronger hydrogen bond donors, and not from extraordinarily precise positioning (*17, 42, 48, 49*). Analogous ensemble–function studies are needed ascertain the origins of oxyanion hole catalysis in other enzymes.

Discussions of dynamics have become nearly inseparable from considerations of enzyme catalysis (e.g., (*12, 26, 55*–*59*)), and the landscape view of catalysis has been presented previously (*7, 11, 12, 60, 61*). Several experimental approaches, including H/D exchange, NMR relaxation, and ensembles from X-ray crystallography provide information about landscapes. Multi-conformer models from room temperature X-ray crystallography provide information about the extent and direction of motion of each atom within a protein and a single additional room temperature crystallographic experiment allows comparison of a new mutant or new liganded state, both of which we took advantage or herein (*26, 34, 52, 62, 63*). Nevertheless, multi-conformer models also have limitations: they do not inform about coupled motions and do not give information about the timescales of motions and interconversions between states. These limitations can be overcome by combining multi-conformer models from room temperature X-ray crystallography with pseudo-ensembles, which can provide information about residues that move in concert (*13, 16, 17*), and H/D exchange and NMR relaxation, which can inform about timescales of motion and interconversion (*64, 65*).

For our KSI mutants, ensemble information from multi-conformer models was needed and sufficient to capture structural effects; double mutant cycles and alternative substrates were needed to identify functional interconnections; and NMR chemical shifts were needed to elucidate changes in active site hydrogen bonds. Approaching mechanism via ensemble–function studies allows cutting edge and vital questions to be addressed, the answers to which are needed to understand the function and evolution of natural enzymes and to design highly efficient enzymes for novel biomedical and industrial applications.

## Materials and Methods

### KSI expression and purification

The ketosteroid isomerase enzymes from *Pseudomonas putida* (referred to herein as KSI, UniProt P07445) was expressed and purified as previously described with minor modifications (*66*). Briefly, BL21 cells transformed with plasmid carrying the desired KSI construct were grown at 37 °C to OD 0.5−0.6 in LB media (EMD Millipore Corp, Billerica, MA, USA) containing 50 μg/mL carbenicillin (Goldbio, St Lousi, MO, USA), and protein expression was induced with 1 mM isopropyl-β-D-1-thiogalactopyranoside (Goldbio, St Lousi, MO, USA). After induction, cultures were grown for 10−12 h at 37 °C. Cells were harvested by centrifugation at 5000 g for 30 min at 4 °C and lysed using sonication. Lysed cells were centrifuged at 48000 g for 30 min at 4 °C. Enzymes were purified from the soluble fraction, first using an affinity column (deoxycholate resin) followed by a size exclusion chromatography column (SEC) Superdex 200. Prior to the purification of each enzyme, the affinity column, FPLC loops, and SEC column were washed with 40 mM potassium phosphate (JT Baker, Omaha, NE, USA), 6 M guanidine (JT Baker, Omaha, NE, USA), pH 7.2 buffer, and then equilibrated with 40 mM potassium phosphate, 1 mM sodium EDTA, 2 mM DTT (Goldbio, St Lousi, MO, USA), pH 7.2 buffer.

### KSI solution kinetics

KSI Michaelis–Menten parameters were obtained by monitoring the 5(10)-estrene-3,17-dione and 5-androstene-3,17-dione (Steraloids, Newport, RI, USA) reaction at 248 nm (extinction coefficient 14,800 M^−1^ cm^−1^) in a PerkinElmer Lambda 25 spectrophotometer (*66*). Reactions were measured at 25 °C in 4 mM sodium phosphate, pH 7.2 buffer with 2% DMSO (JT Baker, Omaha, NE, USA) added for substrate solubility. Low buffer concentrations were used to minimize the background reaction rate. Values of *k*_*cat*_ and *K*_M_ were determined by fitting the initial rates as a function of substrate concentration to the Michaelis–Menten equation. Typically, five to seven substrate concentrations, varying from 2 to 300 μM, were used for each mutant. Averaged values and errors representing the standard deviations are given in Table S2.

### KSI ^1^H solution Nuclear Magnetic Resonance

The ^1^H NMR spectrum of KSI Y57F/D40N bound to (9β,13α)-3-hydroxyestra-1,3,5(10)-trien-17-one was acquired at the Stanford Magnetic Resonance Laboratory using an 800 MHz Varian ^UNITY^INOVA spectrometer running VNMRJ 3.1A and equipped with a Varian 5 mm triple resonance, pulsed field gradient ^1^H[^13^C,^15^N] cold probe, as previously described (*44*). The sample contained 1 mM KSI and 2 mM equilenin (Steraloids, Newport, RI, USA) in 40 mM potassium phosphate (pH 7.2), 1 mM sodium·EDTA, 2 mM DTT, and 10% DMSO-d_6_ (v/v) (Cambridge Isotope Laboratories, Tewksbury, MA, USA). DMSO-d_6_ served as the deuterium lock solvent and prevented freezing at low temperatures. The spectrum was obtained in a 5 mm Shigemi symmetrical microtube at –3.5 °C, following temperature calibration with a 100% methanol standard. The 1331 binomial pulse sequence was used to suppress the water signal with a spectral width of 35 ppm (carrier frequency set on the water resonance) and an excitation maximum between 14-18 ppm (*67*). The data was processed using 10 Hz line broadening and baseline correction applied over the peaks of interest. Chemical shifts were referenced internally to the water resonance.

### Protein crystallization and X-ray data collection

All enzymes were crystallized as previously described (*42*). Briefly, enzyme were crystallized by mixing 1-2 μL of enzyme at 1 mM (for the TSA-bound KSI, preincubated with 2 mM of the TSA (9β,13α)-3-hydroxyestra-1,3,5(10)-trien-17-one) and 1-2 μL, respectively, of crystallization solution (17-23% PEG 3350 (Hampton Research, Aliso Viejo, CA, USA) and 0.2 M MgCl_2_ (JT Baker, Omaha, NE, USA)) in a vapor diffusion hanging drop setup at room temperature. Crystals typically appeared after 24-72 hours. Prior to data collection, crystals were transferred from the crystallization solution to paratone N oil (Hampton Research, Aliso Viejo, CA, USA) where excess crystallization solution was stripped and crystals then were directly mounted on the goniometer for data collection. Because a large body of work identified the 180–220 K temperature range as an inflection point above which both harmonic and anharmonic protein motions are activated, providing strong evidence that protein behavior at 250 K approximates behavior at room temperature (*34*–*36, 68, 69*), and because we previously observed that data collected at 250 K were of slightly higher resolution (∼0.1-0.2 Å) compared to data at 280 K (*17*), we collected data at 250 K. Data collection temperature was controlled using a N_2_ cooler/heater. Single-crystal diffraction data were collected at SSRL, beamline 9-2, using wavelengths of 0.886 Å. See Table S3 for diffraction data statistics.

### Crystallographic data processing and model building

Data processing was carried out with in-house scripts: http://smb.slac.stanford.edu/facilities/software/xds/#autoxds_script. Briefly, data reduction was done using the XDS package (*70*), scaling and merging was done using *Aimless* (*71, 72*) and structure factor amplitudes were obtained using *Truncate* (*71, 73*). Initial phases were obtained via molecular replacement using *PHASER* (*74*) and the PDB entry 3VSY as a search model. Model building was carried out with the program *ARP/wARP* (*75*) and manually in *Coot* (*76*). Traditional, single conformation models, in which major alternative side chain and backbone conformations were modeled, were refined manually after visual inspection with *Coot* and using *phenix*.*refine* (*77*). Torsion-angle simulated annealing (as implemented in *phenix*.*refine*) was used during the initial stages of refinement. Ligand restraints were generated using the *GRADE* server (http://grade.globalphasing.org/cgi-bin/grade/server.cgi). These models were used as input for multi-conformer molding (see below).

Multi-conformer models were obtained using the program *qFit* and previously described methods (*78, 79*). Subsequent to the automated multi-conformer model building, ill-defined water molecules were deleted and alternative protein side and main chain conformations were manually adjusted after visual inspection in *Coot* (*80*) and based on the fit to the electron density. Both alternative side chain rotameric states as well as alternative orientations within the same rotameric state were modeled. Models were subsequently refined with *phenix*.*refine* (*77*). Riding hydrogen atoms were added in the late stages of refinement and their scattering contribution was accounted for in the refinement. Final multi-conformer model quality was checked by *MolProbity* (*81*) and via the PDB Validation server (https://validate-rcsb-2.wwpdb.org/) and deposited on the PDB. See Table S3 for refinement statistics.

### Ensembles vs. multi-conformer models

For both WT and mutants, each KSI state is a dimer and each state is composed of the multi-conformer models for each monomer from the KSI dimer. Thus, the WT ensemble is composed of two multi-conformer models for each of the apo, GSA-bound, and TSA-bound states and we refer to this ensemble of six multi-conformer models as the WT ensemble (*17*). For Y32F/Y57F we obtained multi-conformer models for each of the monomers from the dimer in the apo state. Thus the comparisons in Figures 2, S2, and S5 are made between the WT ensemble made of 6 KSI WT multi-conformer models and the Y32F/Y57F ensemble made of two Y32F/Y57F multi-conformer models. For Y57F we obtained multi-conformer models for each of the monomers from the dimer of both the apo and a TSA-bound states. Thus the comparisons in Figures 2, S2, and S5 are made between the WT ensemble and either the apo Y57F ensemble made of two multi-conformer models (Figure 2) or the TSA-bound ensemble made of two multi-conformer models (Figure 2), or an ensemble made of all four multi-conformer models of Y57F apo and TSA-bound states (Figures S2 and S5).

### Structural alignments

KSI structures and multi-conformer models were aligned on backbone atoms N, CA, C, O of residues 5-125 using PyMOL and standard commands (*82*). The Y32F/Y57F and Y57F multi-conformer models were aligned on the 250 K multi-conformer model of WT apo KSI (PDB 6UCW) as previously described (*17*). The comparison of KSI ensemble or multi-conformer models with traditional single conformation cryo structural models from the PDB (Figures 2 and 4) was achieved by aligning the cryo models on the single conformation model of WT apo KSI that was used to obtain the associated 250 K multi-conformation model in pymol and using standard commands.

## Acknowledgments

Use of the Stanford Synchrotron Radiation Lightsource (SSRL), SLAC National Accelerator Laboratory, is supported by the U.S. Department of Energy, Office of Science, and Office of Basic Energy Sciences under Contract No. DE-AC02-76SF00515. The SSRL Structural Molecular Biology Program is supported by the DOE Office of Biological and Environmental Research and by the National Institute of Health (NIH), National Institute of General Medical Sciences (NIGMS, P41GM103393). The contents of this publication are solely the responsibility of the authors and do not necessarily represent the official views of NIH or NIGMS. This work was funded by a National Science Foundation (NSF) Grant (MCB-1714723) to DH. FY was supported in part by a long-term Human Frontiers Science Program postdoctoral fellowship. MMP and was supported in part by NSF Graduate Research Fellowships and in part by a Lieberman Fellowship (Stanford University). JSF acknowledges funding grant NIH GM123159. We thank Craig Markin for feedback on the manuscript, Michael Thompson for helpful discussion of crystallographic symmetry, and Lisa Dunn (SSRL) for help with scheduling experimental beam time.

