## Supplementary file for "Ensemble–function relationships to evaluate catalysis in the ketosteroid isomerase oxyanion hole"

### Supplementary Information

*Supplementary text 1*

*Hydrogen bond length and energetics.* Proposals to explain enzyme catalysis have invoked a relationship between the hydrogen bond length and the hydrogen bond strength, with the shortest hydrogen bonds proposed to be the strongest (*1–5*). Such short hydrogen bonds are often also called "low barrier" hydrogen bonds which can occur when the hydrogen atom is at the center between the two hydrogen bonding heavy atoms. Low-barrier hydrogen bonds were thought to be unusually strong. These proposals have been discussed extensively and are distinct from hydrogen bond effects addressed herein where hydrogen bond length and energetics arise from reduced polarization of the Y16 KSI hydrogen bond from disruption of the Y57/Y32 hydrogen bond network with Y16 (*1*, *6–13*).

*Supplementary text 2*

*KSI oxyanion hole hydrogen bond length changes*. Because we expect and observe a shortening of the Y16 hydrogen bond with changes in hydrogen bond polarization, there must be some change in the oxyanion hole conformational state. However, for Y32F/Y57F the observed 0.1 Å change in the Y16 hydrogen bond length is modest relative to the conformational motions of Y16 and the oxyanion (on the scale of 1 Å) and therefore not expected to significantly impact the ensemble (*14*). In addition, prior work has established coupling between hydrogen bonds such that shortening of the D103 hydrogen bond is expected to accompany Y16 lengthening, with an effect of 0.3 that of the primary Y16 effect (*11*, *15*). This effect is observed for the Y32F/Y57F (*11*) and Y57F mutants (**Figure S3** and **Table S5**); it is also small relative to overall conformational motions and there are no additional effects of the Y32F/Y57F and Y57F mutations on the conformational states of D103 (see **Figure S2**).

#### Supplementary Figures

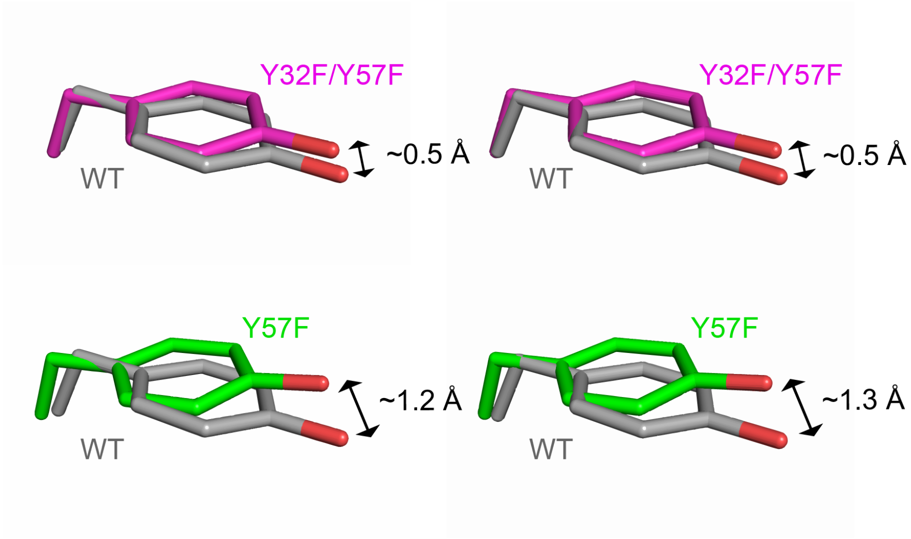

**Figure S1**. Comparison of the Y16 position in WT (PDB 3VSY, grey, (*16*)), Y32F/Y57F (PDB 1DMN, magenta, (*17*)), and Y57F (PDB 1DMM, green, (*17*)). The WT crystal structure contains two molecules (dimer) in the asymmetric unit: molecule A (comparison on the left) and molecule B (comparison on the right), while Y32F/Y57F and Y57F asymmetric unit contains a single KSI molecule (monomer) and the KSI dimer can be reproduced by applying crystallographic two-fold symmetry. The KSI molecules are aligned on the protein backbone (atoms N, CA, C, O, see Materials and Methods). See **Table S1** for RMSDs.

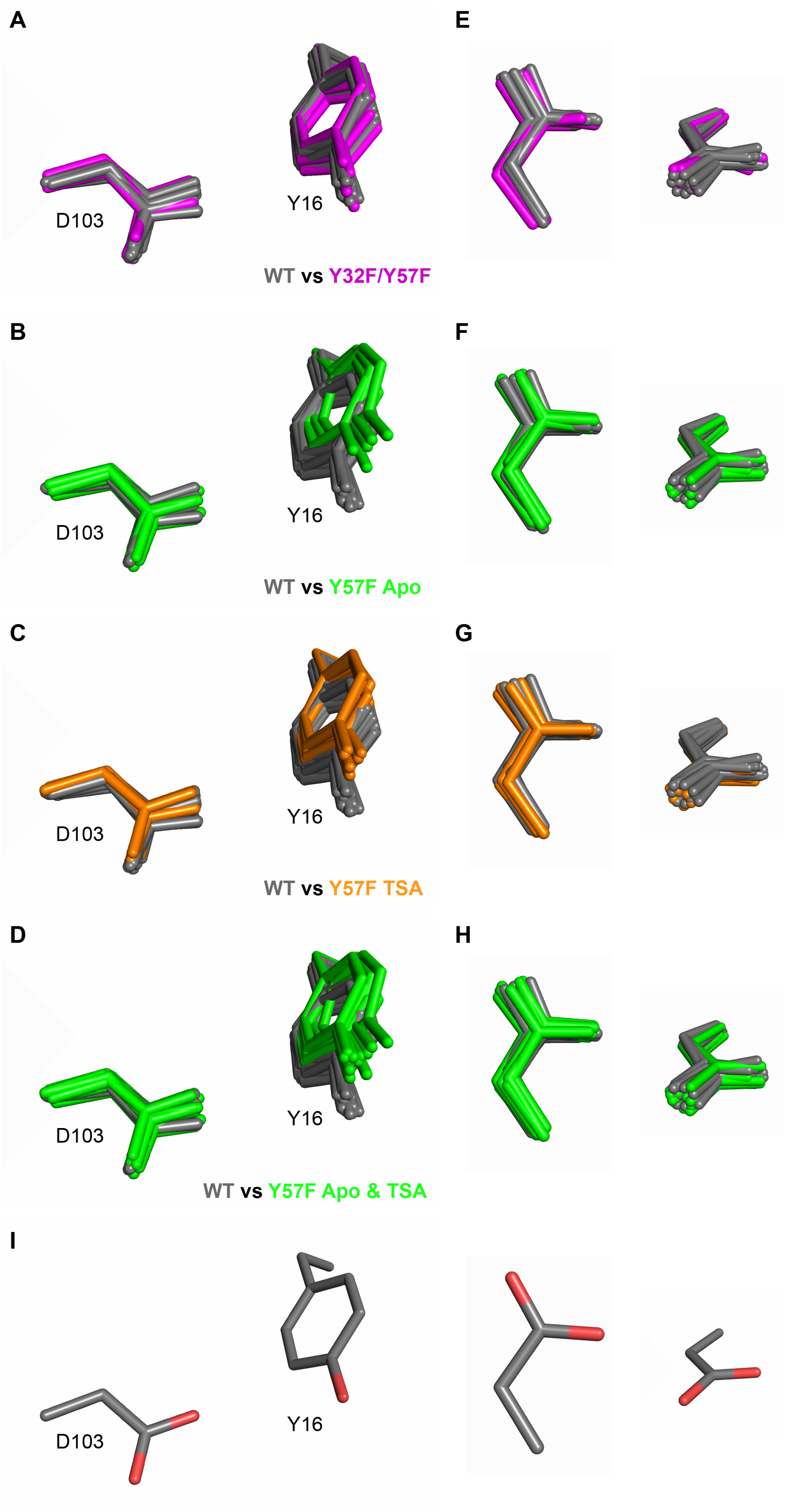

**Figure S2.** Comparison of WT and mutant oxyanion hole Y16 and D103 ensembles and multi-conformer models. Overlay of the WT ensemble (grey) with the multi-conformer models for (**A**) Y32F/Y57F (magenta), (**B**) Y57F apo, (**C**) Y57F TSA-bound (orange), and (**D**) Y57F apo and TSA-bound (green). For both WT and mutants, each KSI state is composed of the multi-conformer models for each monomer from the KSI dimer. Thus, the WT ensemble is composed of 2 multi-conformer models for each of the apo, GSA-bound, and TSA-bound states (*14*). For each mutant, each state is composed of two multi-conformer models. Thus the comparisons in (A-C) are between the WT ensemble and the two Y32F/Y57F apo multi-conformer models, the two Y57F apo multi-conformer models, and the two Y57F TSA-bound multi-conformer models, respectively. In (D) all four Y57F multi-conformer models (apo and TSA-bound) are compared with the WT ensemble. (**I**) Illustrates the orientation of Y16 and D103 in (A-D) but with the oxygen atoms colored in red. (**E-H**) The same comparisons as for (A-D) but now only shown D103 in two different orientations. The comparisons suggest no changes in the D103 ensembles in any of the mutants relative to WT.

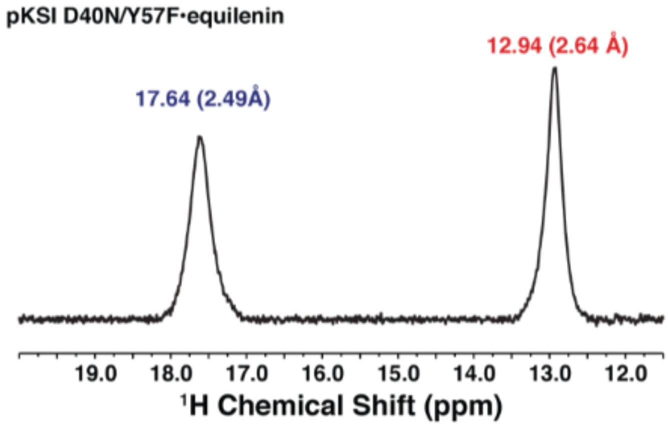

**Figure S3**. $^{1}$H NMR spectrum for Y57F/D40N with bound TSA. The Y57F/D40N – transition state analog complex was prepared and data was collected as described in the Materials and Methods. D40N mutation was introduced to mimic the protonated general base and increase TSA affinity (*9*, *18*).

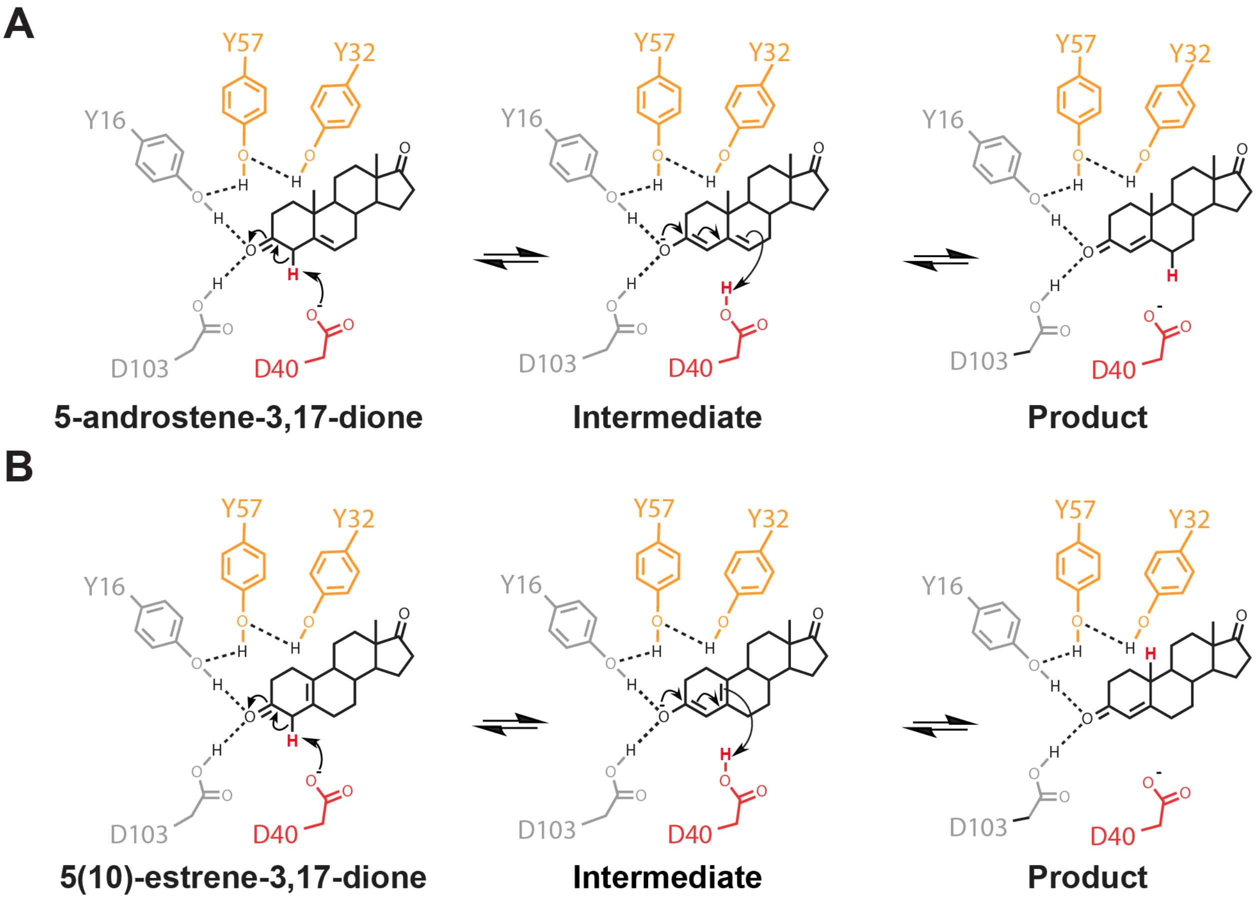

**Figure S4**. KSI reaction mechanism with the steroid substrates 5-androstene-3,17-dione (A) and 5(10)-estrene-3,17-dione (B). The shuffled proton is colored in red.

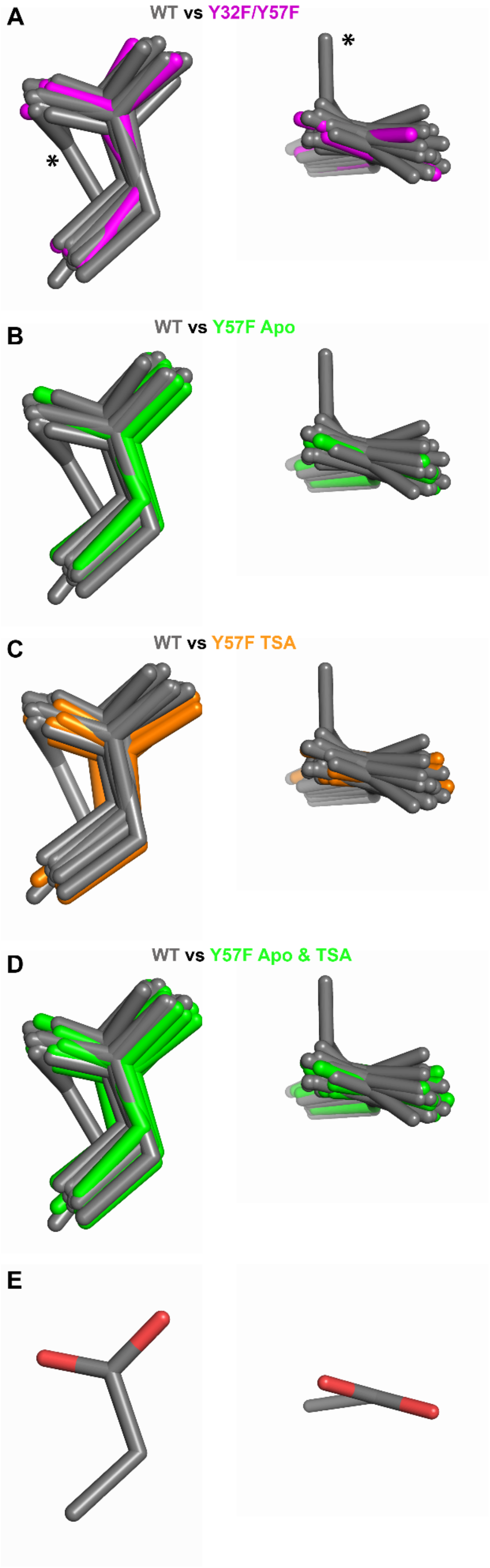

**Figure S5.** Comparison of WT and mutant general base D40 ensembles and multi-conformer models. Overlay of the WT ensemble (grey) with the multi-conformer models for (**A**) Y32F/Y57F (magenta), (**B**) Y57F apo, (**C**) Y57F TSA-bound (orange), and (**D**) Y57F apo and TSA-bound (green). For both WT and mutants, each KSI state is composed of the multi-conformer models for each monomer from the KSI dimer. Thus, the WT ensemble is composed of 2 multi-conformer models for each of the apo, GSA-bound, and TSA-bound states (*14*). For each mutant, each state is composed of two multi-conformer models. Thus the comparisons in (A-C) are between the WT ensemble and the two Y32F/Y57F apo multi-conformer models, the two Y57F apo multi-conformer models, and the two Y57F TSA-bound multi-conformer models, respectively. In (D) all four Y57F multi-conformer models (apo and TSA-bound) are compared with the WT ensemble. (**E**) Illustrates the orientation of general base D40 in (A-D) but with the oxygen atoms colored in red. The asterisk A denotes a general base "out" state that is observed in the complex of WT with bound ground state analog (PDB 6UCY, (*14*)). This seemingly rare "out" state is likely associated with the binding of ground state; its absence within mutant ensembles may be because it remains a rare state or may become less favored; nevertheless, it is not a reactive state and thus not expected to affect absolute or relative reaction rates.

### Supplementary Tables

| RMSD (Å) | 3VSY molecule B | 1DMN | 1DMM |
|---|---|---|---|
| **3VSY molecule A** | 0.15 | 0.26 | 0.26 |
| **3VSY molecule B** | - | 0.25 | 0.25 |

**Table S1.** Alignment of WT (PDB 3VSY, (*16*)), Y32F/Y57F (PDB 1DMN, (*17*)), and Y57F (PDB 1DMM, (*17*)). The WT crystal structure contains two molecules (dimer) in the asymmetric unit, while Y32F/Y57F and Y57F asymmetric unit contains a single KSI molecule (monomer) and the KSI dimer can be reproduced by applying crystallographic two-fold symmetry. The KSI molecules are aligned on the protein backbone (atoms N, CA, C, O, see Materials and Methods).

| Enzyme pKSI | $k_{cat}$ ($s^{-1}$) | $K_M$ (μM) | $k_{cat}$ rel | $k_{cat}$ fold | $K_M$ rel | Reference |
|---|---|---|---|---|---|---|
| **Substrate 5-androstene-3,17-dione** | | | | | | |
| WT | 21230 ± 80 | 49.9 ± 1.3 | (1) | (1) | (1) | (*17*) |
| Y55F | 3510 ± 60 | 23.0 ± 1.0 | 0.17 | 6.0 | 0.5 | (*17*) |
| Y30F/Y55F | 10680 ± 350 | 50.2 ± 5.5 | 0.50 | 2.0 | 1.0 | (*17*) |
| WT | 11000 | 81 | (1) | (1) | (1) | (*19*) |
| Y57F | 1193 ± 220 | 58 ± 26 | 0.11 ± 0.2 | 9.2 ± 1.7 | 0.7 ± 0.3 | This work |
| Y32F/Y57F | 2484 ± 271 | 23 ± 8 | 0.23 ± 0.2 | 4.4 ± 0.48 | 0.3 ± 0.1 | This work |
| **Substrate 5(10)-estrene-3,17-dione** | | | | | | |
| WT | 9.9 ± 0.9 | 30 ± 4 | (1) | (1) | (1) | (*19*) |
| Y57F | 5.05 ± 0.5 | 30.2 ± 6.3 | 0.51 ± 0.1 | 2.0 ± 0.3 | 1.0 ± 0.25 | This work |
| Y32F/Y57F | 2.47 ± 0.3 | 21.9 ± 3.2 | 0.25 ± 0.3 | 4.0 ± 0.6 | 0.7 ± 0.14 | This work |
| D40G | 0.0021 ± 0.0001 | 30 ± 6 | 0.0002 ± 0.00002 | 4.7 x $10^3$ ± 0.5 x $10^3$ | 1.0 ± 0.24 | (*20*) |
| Y57F/D40G | 0.000152 ± 0.00003 | 54.2 ± 5.4 | 0.000015 ± 0.000003 | 65.1 x $10^3$ ± 1.4 x $10^3$ | 1.8 ± 0.30 | This work |
| Y32F/Y57F/D40G | 0.000412 ± 0.000016 | 25.5 ± 6.6 | 0.000042 ± 0.000004 | 24.0 x $10^3$ ± 2.4 x $10^3$ | 0.85 ± 0.25 | This work |
| D40G | 0.0021 ± 0.0001 | 30 ± 6 | (1) | (1) | (1) | (*20*) |
| Y57F/D40G | 0.000152 ± 0.00003 | 54.2 ± 5.4 | 0.07 ± 0.01 | 13.8 ± 2.3 | 0.6 ± 0.1 | This work |
| Y32F/Y57F/D40G | 0.000412 ± 0.000016 | 25.5 ± 6.6 | 0.20 ± 0.01 | 5.1 ± 0.3 | 1.2 ± 0.4 | This work |

**Table S2**. Enzyme kinetics data for KSI WT and mutants with two different substrates: 5-androstene-3,17-dione and 5(10)-estrene-3,17-dione (see Figure S4). The 5-androstene-3,17-dione kinetics data collected in (*19*) and in this work have been used for the analyses carried out. All $K_M$ values are within two fold. Assuming that $K_M$ approximates substrate affinity, the observed 2-fold effects suggest no changes in substrate affinity for any of the mutants.

| | Y32F/Y57F apo | Y57F apo | Y57F TSA-bound |
|---|---|---|---|
| **PDB code** | 7RXK | 7RXF | 7RY4 |
| **Data collection** | | | |
| **Wavelength (Å)** | 0.88557 | 0.88557 | 0.88557 |
| **Resolution range*** | 37.03-1.10 (1.12-1.10) | 35.99-1.16 (1.18-1.16) | 37.28-1.11 (1.13-1.11) |
| **Space group** | $P2_12_12_1$ | $P2_12_12_1$ | $P2_1$ |
| **Unit cell** | 36.27 74.06 96.23 90.00 90.00 90.00 | 35.99 74.14 95.64 90.00 90.00 90.00 | 36.24 74.55 50.97 90.00 110.70 90.00 |
| **Total reflections** | 687514 (31965) | 580641 (26947) | 371916 (17726) |
| **Unique reflections** | 105295 (5087) | 88770 (4193) | 97398 (4738) |
| **Multiplicity** | 6.5 (6.3) | 6.5 (6.4) | 3.8 (3.7) |
| **Completeness (%)** | 99.4 (99.0) | 99.5 (97.8) | 97.8 (95.3) |
| **Mean I/sigma(I)** | 9.2 (0.6) | 7.7 (0.6) | 7.4 (0.7) |
| **R-merge** | 0.079 (2.860) | 0.094 (2.851) | 0.076 (1.764) |
| **R-meas** | 0.086 (3.123) | 0.102 (3.095) | 0.088 (2.051) |
| **R-pim** | 0.033 (1.233) | 0.039 (1.192) | 0.044 (1.028) |
| **$CC_{1/2}$**** | 0.999 (0.315) | 0.998 (0.435) | 0.999 (0.303) |
| **Refinement** | | | |
| **Resolution range** | 34.56-1.10 (1.14-1.10) | 32.38-1.16 (1.20-1.16) | 29.37-1.11 (1.15-1.11) |
| **Reflections used in refinement*** | 104743 (10013) | 88211 (8300) | 97166 (9355) |
| **Reflections used for R-free** | 5233 (472) | 4393 (413) | 4893 (498) |
| **R-work** | 0.153 (0.337) | 0.152 (0.337) | 0.143 (0.300) |
| **R-free** | 0.166 (0.324) | 0.171 (0.342) | 0.158 (0.300) |
| **non-hydrogen atoms** | 5199 | 5241 | 6277 |
| **macromolecules** | 4870 | 4985 | 5850 |
| **ligands** | 6 | 4 | 130 |
| **solvent** | 323 | 252 | 297 |
| **Protein residues** | 257 | 254 | 257 |
| **RMS(bonds)** | 0.007 | 0.009 | 0.008 |
| **RMS(angles)** | 0.95 | 1.02 | 1.07 |
| **Ramachandran favored (%)** | 97.6 | 97.2 | 96.8 |
| **Ramachandran allowed (%)** | 2.4 | 2.8 | 3.2 |
| **Ramachandran outliers (%)** | 0.0 | 0.0 | 0.0 |
| **Average B-factor** | 17.8 | 19.9 | 14.5 |
| **macromolecules** | 17.0 | 19.3 | 13.8 |
| **ligands** | 17.0 | 21.5 | 16.6 |
| **solvent** | 29.4 | 31.6 | 27.1 |

*Values in parenthesis are for the highest resolution shells. ** $CC_{1/2}$ values are for the following resolution shells 7RXK: 1.10-1.13 Å, 7RXF: 1.16-1.19 Å, 7RY4: 1.11-1.14 Å.

**Table S3.** X-ray diffraction and model refinement statistic. All data collection statistics were obtained from Aimless (*21*, *22*), with the exception of $CC_{1/2}$, which was obtained from XSCALE from the XDS package (*23*). Model refinement statistics were obtained from phenix (*phenix.table_one*) using the final refined models and reflections files deposited on the PDB.

| RMSD (Å) | WT Apo molA | WT Apo molB | Y32F/Y57F Apo molA | Y32F/Y57F Apo molB | Y57F Apo molA | Y57F Apo molB | Y57F TSA molA | Y57F TSA molB |
|---|---|---|---|---|---|---|---|---|
| **WT Apo molA** | - | 0.31 | 0.25 | 0.45 | 0.35 | 0.40 | 0.39 | 0.41 |
| **WT Apo molB** | 0.31 | - | 0.35 | 0.35 | 0.41 | 0.42 | 0.42 | 0.40 |

**Table S4.** Alignment of multi-conformer models for WT and mutants. WT and mutant KSI crystals contain two molecules (dimer) in the asymmetric unit (molecule A and molecule B). RMSDs were calculated for each KSI molecule independently via alignment on the protein backbone (atoms N, CA, C, O, see Materials and Methods).

| Enzyme | ΔHB length (Y16) Å | Reference |
|---|---|---|
| WT | (0) | (*11*) |
| Y32F | 0.025 | (*11*) |
| Y32F/Y57F | 0.1 | (*11*) |
| D103N | 0.17 | (*11*) |
| Y57F | 0.14 | This work |

**Table S5.** Hydrogen bond length changes (ΔHB length) were obtained from $^1$H solution NMR as previously described ((*9*, *11*) and see Materials and Methods).

| Enzyme | $k_{cat, rel}$ | $k_{cat}$ rel | Log $k_{cat, rel}$ | ΔHB length (Y16) Å |
|---|---|---|---|---|
| WT | 11000 | (1) | 0 | 0 |
| Y32F | - | - | - | 0.025 |
| Y32F/Y57F | 2484 | 0.23 | -0.64 | 0.1 |
| Y57F | 1193 | 0.11 | -0.96 | 0.14 |
| D103N | - | - | - | 0.17 |

**Table S6**. Enzyme kinetics data for KSI WT and mutants with substrate 5-androstene-3,17-dione (from Table S2) and changes in hydrogen bond length (ΔHB length, from Table S5) used in Figures 2F and 3E.

| Enzyme | $k_{cat}$ | $k_{cat, rel}$ | Log $k_{cat, rel}$ | Enzyme | $k_{cat}$ | $k_{cat, rel}$ | Log $k_{cat}$, rel | ΔHB length (Y16) Å |
|---|---|---|---|---|---|---|---|---|
| WT | 9.9 | (1) | 0 | D40G | 0.0021 | (1) | 0 | 0 |
| Y32F | 5 | 0.5 | -0.3 | Y32F/ D40G | - | - | - | 0.025 |
| Y32F/ Y57F | 2.5 | 0.25 | -0.6 | Y32F/ Y57F/ D40G | 0.00041 | 0.195 | -0.71 | 0.1 |
| Y57F | 5.1 | 0.51 | -0.29 | Y57F/ D40G | 0.000152 | 0.072 | -1.14 | 0.14 |
| D103N | 0.76 | 0.077 | -1.11 | D103N/ D40G | - | - | - | 0.17 |

**Table S7.** Enzyme kinetics data for KSI WT and mutants relative to WT (D40 background) or relative to D40G with substrate 5(10)-estrene-3,17-dione (from Table S2) and changes in hydrogen bond length (ΔHB length, from Table S5) used in Figures 2F and 3E.
